# Diffusion on PCA-UMAP manifold captures a well-balance of local, global, and continuum structure to denoise single-cell RNA sequencing data

**DOI:** 10.1101/2022.06.09.495525

**Authors:** Cristian Padron-Manrique, Aarón Vázquez-Jiménez, Diego Armando Esquivel-Hernandez, Yoscelina Estrella Martinez Lopez, Daniel Neri-Rosario, Jean Paul Sánchez-Castañeda, David Giron-Villalobos, Osbaldo Resendis-Antonio

## Abstract

Single-cell transcriptomics (scRNA-seq) is becoming a technology that is transforming biological discovery in many fields of medicine. Despite its impact in many areas, scRNASeq is technologically and experimentally limited by the inefficient transcript capture and the high rise of noise sources. For that reason, imputation methods were designed to denoise and recover missing values. Many imputation methods (e.g., neighbor averaging or graph diffusion) rely on k nearest neighbor graph construction derived from a mathematical space as a low-dimensional manifold. Nevertheless, the construction of mathematical spaces could be misleading the representation of densities of the distinct cell phenotypes due to the negative effects of the curse of dimensionality. In this work, we demonstrated that the imputation of data through diffusion approach on PCA space favor over-smoothing when increases the dimension of PCA and the diffusion parameters, such *k-NN* (*k-nearest neighbors*) and *t* (value of the exponentiation of the Markov matrix) parameters. In this case, the diffusion on PCA space distorts the cell neighborhood captured in the Markovian matrix creating an artifact by connecting densities of distinct cell phenotypes, even though these are not related phenotypically. In this situation, over-smoothing of data is due to the fact of shared information among spurious cell neighbors. Therefore, it can not account for more information on the variability (from principal components) or nearest neighbors for a well construction of a cell-neighborhood. To solve above mentioned issues, we propose a new approach called sc-PHENIX( ***s**ingle **c**ell-**PHE**notype recovery by **N**on-linear **I**mputation of gene e**X**pression*) which uses PCA-UMAP initialization for revealing new insights into the recovered gene expression that are masked by diffusion on PCA space. sc-PHENIX is an open free algorithm whose code and some examples are shown at https://github.com/resendislab/sc-PHENIX.

## Introduction

Single-cell RNA sequencing (scRNA-seq) is a revolutionary technique that simultaneously reveals the transcriptome profiles of thousands of individual cells disaggregated from human tissue or cultured cells in vitro. Thus, it provides a higher biological resolution than bulk RNA-sequencing proper to explore questions around the structural and functional heterogeneity inside and among biological samples [1].

However, scRNA-seq protocols suffer from various noise sources, the most detrimental effect is the ‘dropout events’, where a gene has low or high-level expression in one cell but is not detected in another cell of the same cell type [2]. The effect of these events is reflected in the count matrix of a typical scRNA-Seq experiment, with an excess of zeros and only a small fraction of transcripts (~10% – 15%) detected in each cell [3,4]. Therefore, due to this inherent stochasticity, produced by the low initial concentration of mRNA for individual cells, gene-gene relationships profiles are lost and only the strongest relationships between genes prevail [5].

With the purpose to recover the missing expression values and reducing systematic technical noises, imputation strategies combined with dimensional reduction methods have been suggested [6]. These methods have significantly contributed to extracting meaningful biological information and identifying novel cell subpopulations [7].

The task of reducing the dimensionality in scRNA-seq data is difficult because euclidean distances among single-cells tend to be homogeneous in high-dimensional space(‘*curse of dimensionality’*) [8]. Consequently, several samples from distinct single-cell phenotypes could become spurious nearest-neighbors in the low dimensional representation of these methods[8,9]. Different dimensional reduction methods approaches have been proposed to deal with the high-dimensionality of scRNA-seq data like PCA [10], t-SNE [11], and UMAP [12]. All reductional methods focus on preserving local, global, or continuum structure or a balance of these structures. In mathematical spaces (here, embeddings through dimensionality reduction), the global structure only considers the arrangement of distinct clusters, rendering an overall view of the system (typically resulting in a noisy visualization where distinct clusters are overlapped) [13]. The local structure reveals fine-grained details within inner clusters and is often seen as a defined separation of distinct clusters(relevant information of the heterogeneity). Continuum structure is often described as trajectories or branches, samples go through a continuum of gradual and progressive changes (in our case, changes in transcriptomic single-cell profiles) [13]. Here, we consider that the well-balance of a global and local structure permits a well-continuum structure among true connected but distinct clusters. a real biological transition among distinct but transition-involved cell phenotypes).

Although reductional methods have particular advantages and limitations. For instance, PCA tends to give a more faithful representation of the global structure of data [13]. However, the local structure is lost and it is difficult to recognize the fine-grained structure. Regarding non-linear visualization methods such as UMAP and t-SNE, they often scramble the data global structure. Moreover, they are well-suited to recognize the fine-grained local structure of the scRNA-seq data. UMAP arguably preserves more of the global structure and it has a faster time performance than t-SNE [12]. In addition using PCA as an informative initialization step for UMAP or t-SNE helps to preserve the global data structure [14]. Moreover, it has been shown that PCA-UMAP dimensionality reduction reveals fine-scale population structure and is computationally more advantageous compared to PCA-tSNE [15]. However, PCA, t-SNE, and UMAP are not designed to capture a well-continuum structure [13]. For example in visual embeddings, t-SNE and UMAP tend to break data into discrete clusters even when data has trajectories. Unfortunately, t-SNE can only reduce dimensionality to fewer than or equal to 3 dimensions, leading the information of high-dimensional data to get lost. In contrast, UMAP has no such methodological restrictions on embedding dimensions and gets closer to the dimension of the underlying manifold on which sc-RNA-seq data lie. The embedding obtained from these reductional methods has been successfully used on methods to analyze and find functional relations among scRNA-seq data [6,13,14,16,17], one example is imputation [6].

Concerning scRNA-seq data imputation, the imputation methods are capable of recovering the expression matrix to reflect a more accurate transcriptome scenario; they fall into the following categories: model-based, smoothing-based, machine learning-based, and matrix theory-based methods. The different scRNA-seq imputation methods have already been benchmarked elsewhere [6,18]. MAGIC and SAVER have better performance than other methods given their capacity to identify differentially expressed genes, unsupervised clustering, trajectory analyses, memory usage, and scalability, among other evaluations [5,19]. SAVER (model-based method) imputes data by modeling the observed counts as a negative binomial distribution (Poisson-Gamma mixture). Here, technical noise in the gene expression signal is approximated by the Poisson distribution, while the gamma prior distribution accounts for the uncertainty in the true expression. The recovered expression is a weighted average of the normalized observed counts and the predicted true counts [19]. MAGIC (smoothing-based method) imputes by sharing information across similar cells via data diffusion. Thus, it is based on the diffusion maps theory [20]. Firstly, it computes a distance matrix of cells on PCA space, and an affinity matrix is built by applying an adaptive kernel to the distance matrix. Then, a diffusion process is applied to denoise the affinity matrix. Lastly, to recover gene expression, the original data is multiplied by the denoised affinity matrix [5]. Although SAVER and MAGIC recover lost data in many cases, a drawback of these imputation methodologies is the false association (“over-smoothing”) of the data resulting in the consideration of inaccurate gene–gene interactions [21].

In this context, it is clear the need to develop new imputation methods that include the advantages of the state of the art methods and improve the performance in gene expression imputation. To contribute in this field, we developed sc-PHENIX (***s**ingle **c**ell-**PHE**notype recovery by **N**on-linear **I**mputation of gene e**X**pression*) an open-source Python package that implements an hybrid unsupervised machine learning approach for recovering missing gene expression in single-cell data (Fig 1). We found that sc-PHENIX applying a diffusion process on the PCA-UMAP space (PCA-UMAP space as initialization) solves the major issues that come from diffusion on PCA space.

**Fig 1.**
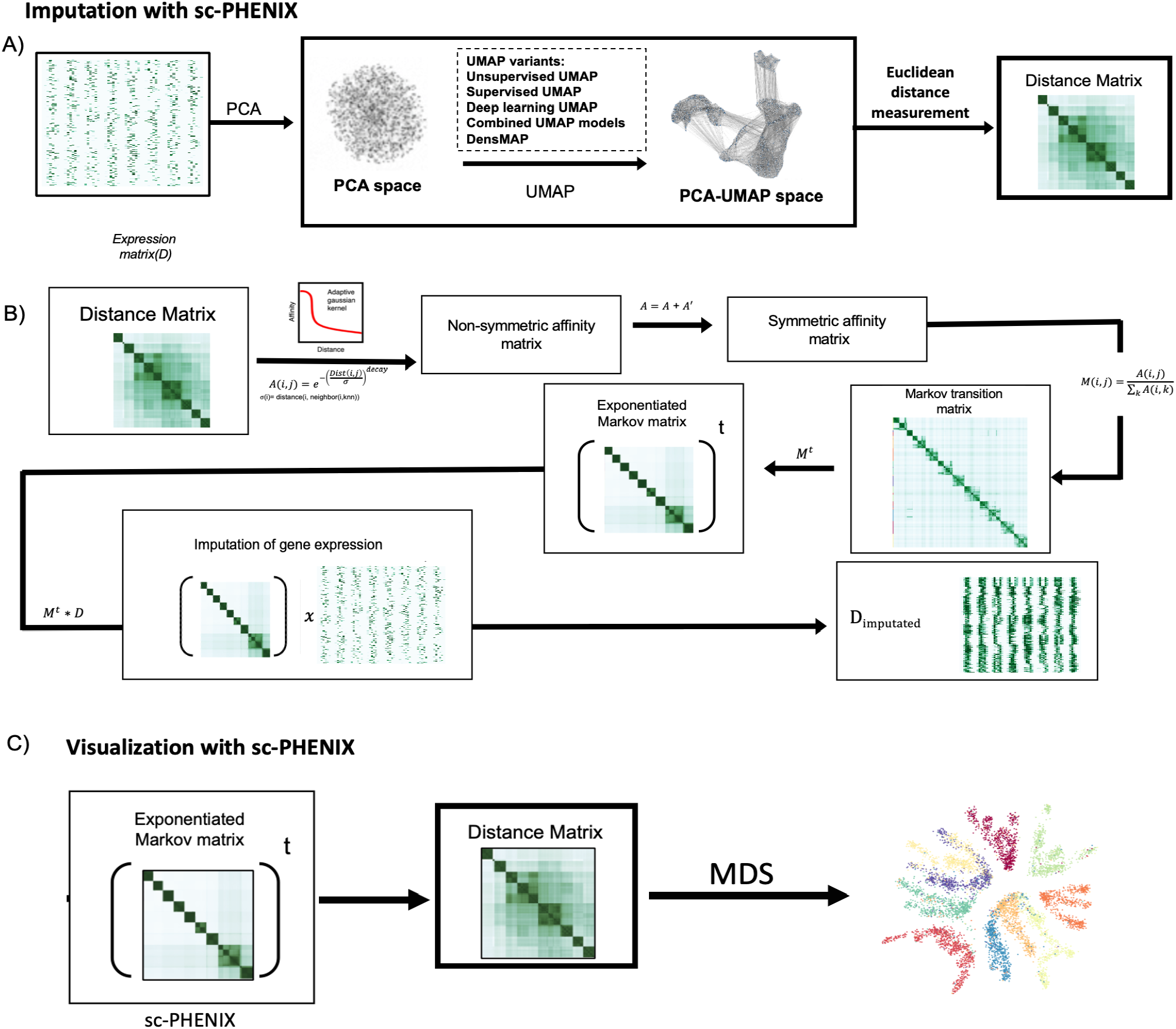
The imputation process using sc-PHENIX. The sc-PHENIX imputation approach for scRNA-seq data consists of two main steps A) The construction of the distant matrix by applying different strategies of linear and non-linear dimensionality reduction, such as PCA or UMAP respectively. sc-PHENIX is characterized by applying consecutively PCA then UMAP. In this PCA-UMAP multidimensional space, sc-PHENIX constructs the best denoise representation of cell distances measurement. B) The diffusion maps for imputation: the imputation process using diffusion maps consist of several steps: 1. Construction of Markov transition matrix from the distant matrix; sc-PHENIX uses the adaptive kernel to generate a non-symmetric affinity matrix, it is symmetrized and then is normalized to generate the Markov transition matrix. 2. Diffusion process; the Markov transition matrix *M* is exponentiated to a chosen power *t* (random walk of length *t* named “diffusion time”). 3. Imputation; imputation of gene expression consists in multiplying the exponentiated Markov matrix (*M^t^*) by the single cell-matrix data to obtain an imputed and denoised single-cell RNA-seq matrix. Equations are described in (**Methods)**. C) Visualization of the exponentiated Markov matrix. For that, we convert the *M^t^* in a distance matrix. Then we apply a multidimensional scaling method to project data in 2D or 3D dimensions. This projection can be used as a heuristic method for quality control of imputation.

## RESULTS

### The sc-PHENIX method

This software was developed to improve imputation of sparse data avoiding over-smoothing. Therefore, our method captures local, global, and continuum structure of data. There is always some inherent danger in over-smoothing the data by running dimensional reduction techniques like PCA, after having performed smoothing/imputation. For that reason, to improve the imputation of data, we consecutively applied PCA and UMAP on the count matrix, then we build a distant matrix through an euclidean metric between all the pairs of data in a multidimensional space. In this PCA-UMAP multidimensional space, sc-PHENIX represents data into a reduced projection which generates the best denoise representation of cell distances measurement for the diffusion process. Having computed the distance matrix, we constructed the Markov transition matrix by applying a similar approach in MAGIC [14]. After that, we applied the diffusion process by exponentiating the transition Markov matrix (*M*) to a chosen power *t* (random walk of length t named “diffusion time”). At this point, the imputed and denoised scRNA-seq matrix is obtained by multiplying the single cell-matrix data with the exponentiated Markov transition matrix (*M^t^* The algorithm is described in more detail in Fig 1 and in the methods section. Finally to have a quality control of imputation, we also provide a diagnostic plot to visualize the *M^t^* cell-neighborhood. For example, the gene expression information will be shared incorrectly among distinct cell phenotypes if they are the spurious nearest neighbors in this plot.

Our method falls into the category of smooth-based imputation. However, the methods used in sc-PHENIX to obtain the low representation manifold (UMAP) and the *M^t^* (diffusion maps) are techniques based on Manifold learning (nonlinear dimensionality reduction methods, a subfield of machine learning) such as diffusion maps and UMAP. The method sc-PHENIX follows the same assumption in manifold learning that the observed data lie on a low-dimensional manifold embedded in a higher-dimensional space.

### Visualization of the exponentiated Markov matrix based on different manifolds

In sc-PHENIX, or any general diffusion-based method, the diffusion process occurs when the transition Markov matrix (*M*) is exponentiated, Fig 1B. The exponentiated transition Markov matrix (*M^t^*) step is essentially a low-pass filter that increases the weighted affinities for similar data, whereas spurious neighbors are down-weighted [14]. The *M^t^* is a weighted graph that shows the probabilities of transitioning among samples (a *single-cell* sample in case of scRNA-seq data) in the data using random walks of any (*t*) length, rendering a temporal ordering of samples [20]. The information provided by the *M^t^* has been used to impute data via diffusion on PCA space, sharing information through local neighbors that follow data continuum densities in a process that is mathematically akin to diffusing heat through the data [5]. To analyze the sample/cell neighborhood captured by the *M^t^*, we compare the *M^t^* matrix constructed by MAGIC to the *M^t^* of sc-PHENIX (using diffusion PCA-UMAP space). However MAGIC returns only the imputed matrix, not its *M^t^*. For that purpose, we indirectly calculate the *M^t^* of MAGIC by computing the *M^t^* with sc-PHENIX with PCA initialization. (See section S1 to further details about the reproducibility of sc-PHENIX as MAGIC with PCA space).

We found that applying diffusion on the PCA space (computed as MAGIC) distorts the underlying manifold of the data by connecting near regions between the densities of distinct phenotypes. Thus, the *M^t^* increases the similarity among densities of distinct cell phenotypes enhancing the over-smoothing in data imputation via diffusion. To observe the distortion effect of diffusion on different manifolds (represented in the *M^t^*), we compared the data similarities using the multidimensional scaling (MDS) projections. On the MDS the distance magnitude is inversely correlated to the similarity. So, we computed the distance matrix taking the *M^t^* from PCA space, and PCA-UMAP space (our contribution) of the same data. For each distance matrix a MDS plot was generated. In addition, to evaluate the method capabilities we used two different datasets: the MNIST database of handwritten digits [23], and the adult mouse visual cortex cells scRNA-seq [24] datasets (Fig 2).

**Fig 2.**
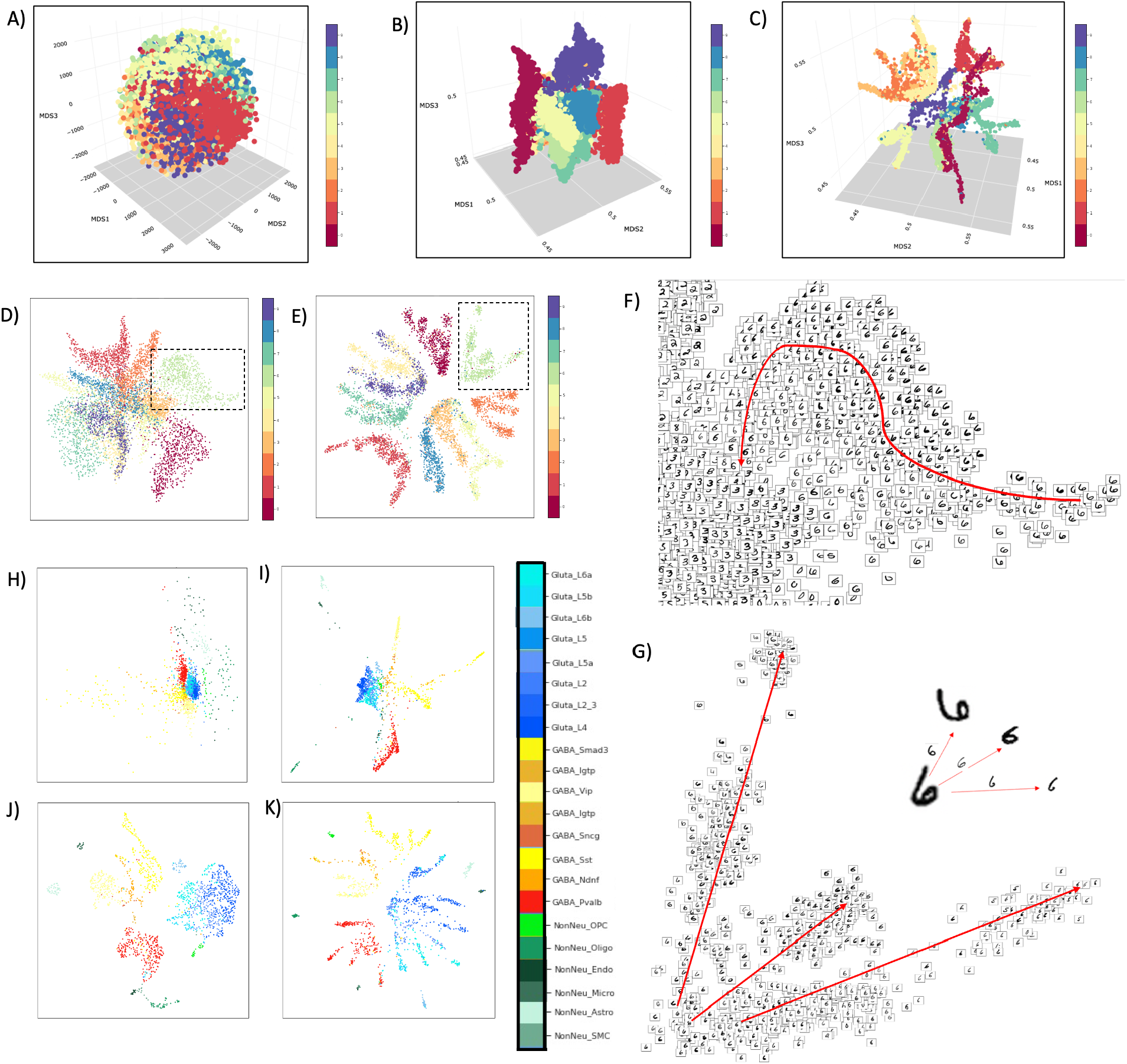
Multidimensional scaling visualization of the exponentiated Markov matrix of sc-PHENIX. The PCA and PCA-UMAP manifold of the MNIST digit and the scRNA-seq of the adult mouse visual cortex cells dataset was used as input for sc-PHENIX to get the exponentiated Markov matrix. The exponentiated Markov matrix was transformed into a distance matrix to be visualized by an MDS process. A. 3D MDS plot of the PCA manifold (500 PC’s, MNIST). B. 3D MDS plot of the exponentiated Markov matrix (500 PC’s input, MNIST). C. 3D MDS plot of the exponentiated Markov matrix (500 PC’s transformed into 60 UMAP components as input, MNIST). D. 2D MDS plot of the exponentiated Markov matrix (500 PC’s input, MNIST). E. 2D MDS plot of the exponentiated Markov matrix (500PC’s transformed into 60 UMAP components as input, MNIST). F. One branch of the 6’s digit images of the PCA space (subsection of Fig 2 D). Redline color indicates the branch continuum G. Three branches of the 6’s digit images subsection of the PCA-UMAP (subsection of the Fig 2 E). H. 2D MDS plot of the PCA manifold (500 PC’s input, scRNA-seq data of the adult mouse visual cortex cells dataset. I. 2D MDS plot of the exponentiated Markov matrix (500PC’s input scRNA-seq data) of the adult mouse visual cortex cells dataset. J. 2D UMAP plot (scRNA-seq data) of the adult mouse visual cortex cells dataset. K. 2D MDS plot of the exponentiated Markov matrix (500PC’s transformed into 60 UMAP components as input, scRNA-seq data) of the adult mouse visual cortex cells dataset. Note: For the adult mouse visual cortex cells dataset, there are three main clusters are GABAergic (red-yellow-ish), glutamatergic (blue-ish) and non-neuronal (green-ish) cell types.

In the MDS plots ( described in more detail in the methods section) of the PCA space in both datasets, we observed distinct digit number images (Fig 2A and Fig S1) and distinct cell phenotype densities (Fig 2H) are close to each other on the PCA space, only separated by a small breach (Fig S2). This is because of the undesired effects of distance concentration (distances among samples in high-dimensionality tend to be closer) [4]. Therefore, the global structure is more preserved than the local structure, a feature of PCA.

On the other hand, by applying the diffusion process and visualizing the *M^t^* constructed from PCA space, reveals some fine-grained local structure in both datasets, MNIST (Fig 2B and D, and Fig S2) and the scRNA-seq (Fig 2I). However, as one can see in Fig 2B, D, I and Fig S2 the local structure gets lost due to the connection of the near regions between the densities of different clusters. As a consequence of the local structure lost, there is a not defined continuum structure.

For example, we can see in Fig 2F and Fig S8 (diffusion on PCA space) that the number 6 images change their shape gradually through the data continuum in a single branch. However, the “6” digit cluster density is connected with all digit cluster densities. In contrast, in Fig 2G and Fig S8 (diffusion on PCA-UMAP space), the “6” digit cluster shows more branches and is separated from distinct digit clusters (Fig 2C, E, and Fig S8). Thus, it indicates that diffusion on PCA-UMAP captures a better representation of the local and the continuum structure. A similar conclusion is obtained for the sc-RNAseq data, for diffusion on PCA (Fig 2I) and diffusion on PCA-UMAP space (Fig 2K).

Previous results showed the spurious connection of different clusters located in the manifold’s center caused by diffusion on PCA space In MNIST (Fig. 2B, D, and Fig S8) could be due to handwritten numbers images having similar shapes in the near regions between the densities of distinct digit clusters. However, there are no similar shapes among digit images in the center, Fig S8.

In sc-RNA seq data (Fig 2I, diffusion on PCA space), the interpretation of the nearness of the different clusters (distinct phenotypes) could indicate that different clusters (different cell phenotypes) are similar biologically/phenotypically. However, in biological reality they are not, GABAergic and glutamatergic neuronal cell phenotypes carry out divergent neuronal functions [24]. Thus, they could not be nearest neighbors (similar phenotypes) in Fig 2I manifold. Moreover, some non-neuronal phenotypes are nearest neighbors of the neuronal cells in Fig 2I (green-ish colors).

Furthermore, we quantified the distribution of MNIST images to detected dense regions (Fig 3). The idea of this analysis is that dense regions indicate an agglomeration of similar MNIST images when an accurate approximation of the underlying manifold is obtained. For this, we use HDBSCAN (*Hierarchical Density-Based Spatial Clustering of Applications with Noise*)[16] on the distance matrix from PCA space, and each of the *M^t^* constructed from PCA space and PCA-UMAP space. HDBSCAN associates as noise samples, samples that do not belong to a dense region. We found that in PCA space, one dense region and a small cluster. This cluster has only a few number “1” of the MNIST samples and most MNIST samples are considered noise (Fig 3A and 3D). In diffusion on PCA space, there is one dense cluster detected (Fig 3B and 3E, Cluster A). This cluster has almost all MNIST samples. However, a few MNIST samples are considered noise (Fig 3B and 3E) compared to the PCA manifold (Fig 3A). In diffusion on PCA-UMAP space, the distribution of densities of distinct MNIST images are well separated in 7 clusters (A-G clusters), see Fig 3C and 3F.

**Fig 3.**
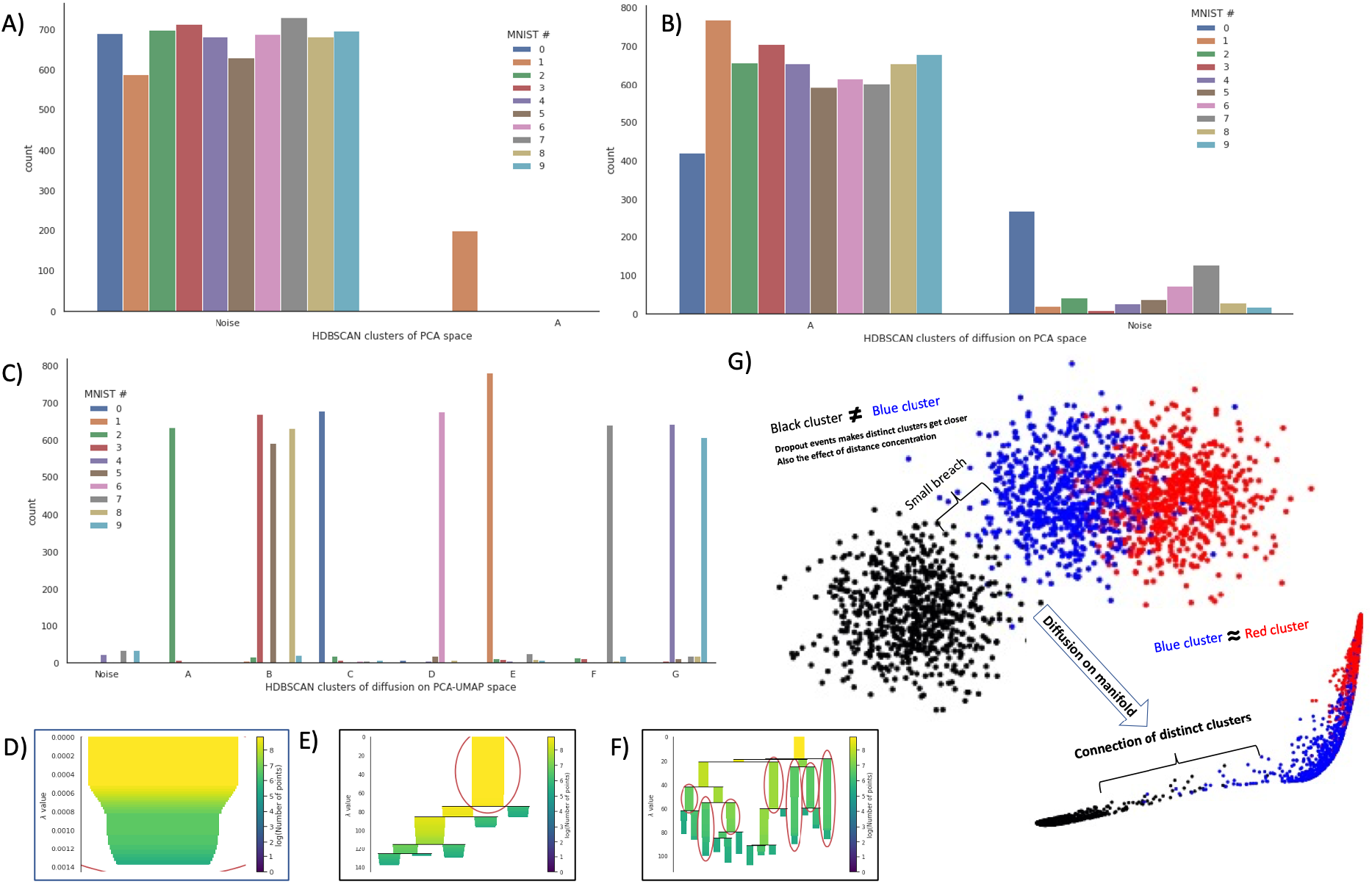
HDBSCAN clusters on the exponentiated Markov matrix of sc-PHENIX. We use HDBSCAN (Hierarchical Density-Based Spatial Clustering of Applications with Noise) on the distance matrix from PCA space, and each of the Mt constructed from PCA space and PCA-UMAP space. HDBSCAN determines noise to samples that do not belong to a dense region. Here, clusters were assigned as letters (for example A,B,C,.etc). A) MNIST samples distribution of different HDBSCAN clusters (PCA space). B) Distribution of MNIST samples on different HDBSCAN clusters of the *M^t^*(diffusion on PCA space). C) Distribution of MNIST samples on different HDBSCAN clusters of the *M^t^*(diffusion on PCA-UMAP space) D) Condense tree plot (PCA space). E) Condense tree plot (diffusion on PCA space). F) Condense tree plot (diffusion on PCA-UMAP space). G) Schematic representation of the diffusion process on lower dimensional representation affected on a manifold representation that is of high dimensionality. We can see that the manifold (in this case the PCA space) by applying a diffusion on, makes the two different clusters (black and blue) connect.This connection occurs in the close regions between the different clusters, separated by a small space. Therefore, after the diffusion process, an artifact consisting of spurious neighboring samples that do not actually have similar features.

Additionally, we wanted to see the difference of capturing local and continuum structure of the data between our visualization method (MDS plots of the *M^t^* from diffusion on PCA-UMAP space of sc-PHENIX) and PHATE (a specialized visualization method). PHATE is a diffusion based method that was designed to preserve local, global and continuum structures in the data for visualization purposes [13]. PHATE uses PCA to improve the robustness and reliability of their *M^t^*, also MAGIC. However, in PHATE plots of the neuronal dataset (Fig S6 and Fig S7), non-neuronal cells (green cluster) are overlapped with neuronal phenotype clusters. A data continuum of cells is present but local structure is lost by overlapping some non-related clusters. The same above conclusion with PHATE happens on MNIST data (Fig S4 and S5). In contrast to PHATE, UMAP is not well designed to preserve continuum structure in the data [13]; it preserves discrete cluster densities, even when the dataset is connected, see UMAP plot (Fig 2J). However, if diffusion is applied on multidimensional PCA-UMAP space(Fig 2K), the local and continuum structure is well captured (as sc-PHENIX using PCA-UMAP space).

Therefore, all indicate that the connection between densities of distinct clusters (images or cell phenotypes) is an artifact (defect) by diffusion on a not well-suited manifold initialization (such as PCA space). This artifact is driven by a small breach between densities of distinct clusters in PCA space (depicted in Fig 3G). Here we show that PCA does not separate well densities from distinct clusters enough to avoid spurious connections by diffusion (Fig 3G). The most probable reason is that sparsity has a major impact on the nearness of densities from distinct clusters in PCA space (Section 2). However, according to our results PCA-UMAP initialization solves this major issue for the diffusion process.

Last but not least, for imputation, sc-PHENIX method requires the multiplication of *M^t^* by the count matrix. To avoid over-smoothing, an effect caused by sharing information among spurious nearest-neighbors, the *M^t^* needs to capture the local and continuum structure of the data as well as possible (global structure to some extent). For that reason, we recommend sc-PHENIX using PCA-UMAP(Fig 2K) space instead of only PCA to solve the major issues caused by diffusion on PCA space(Fig 2I).

### Evaluation of Peripheral Blood Mononuclear Cells Dataset

To continue with the sc-PHENIX evaluation and test their applicability, we used the dataset of 3k Peripheral Blood Mononuclear Cells (PBMC) freely available from 10X Genomics. This dataset has different cell phenotypes(Fig 4).

**Fig 4.**
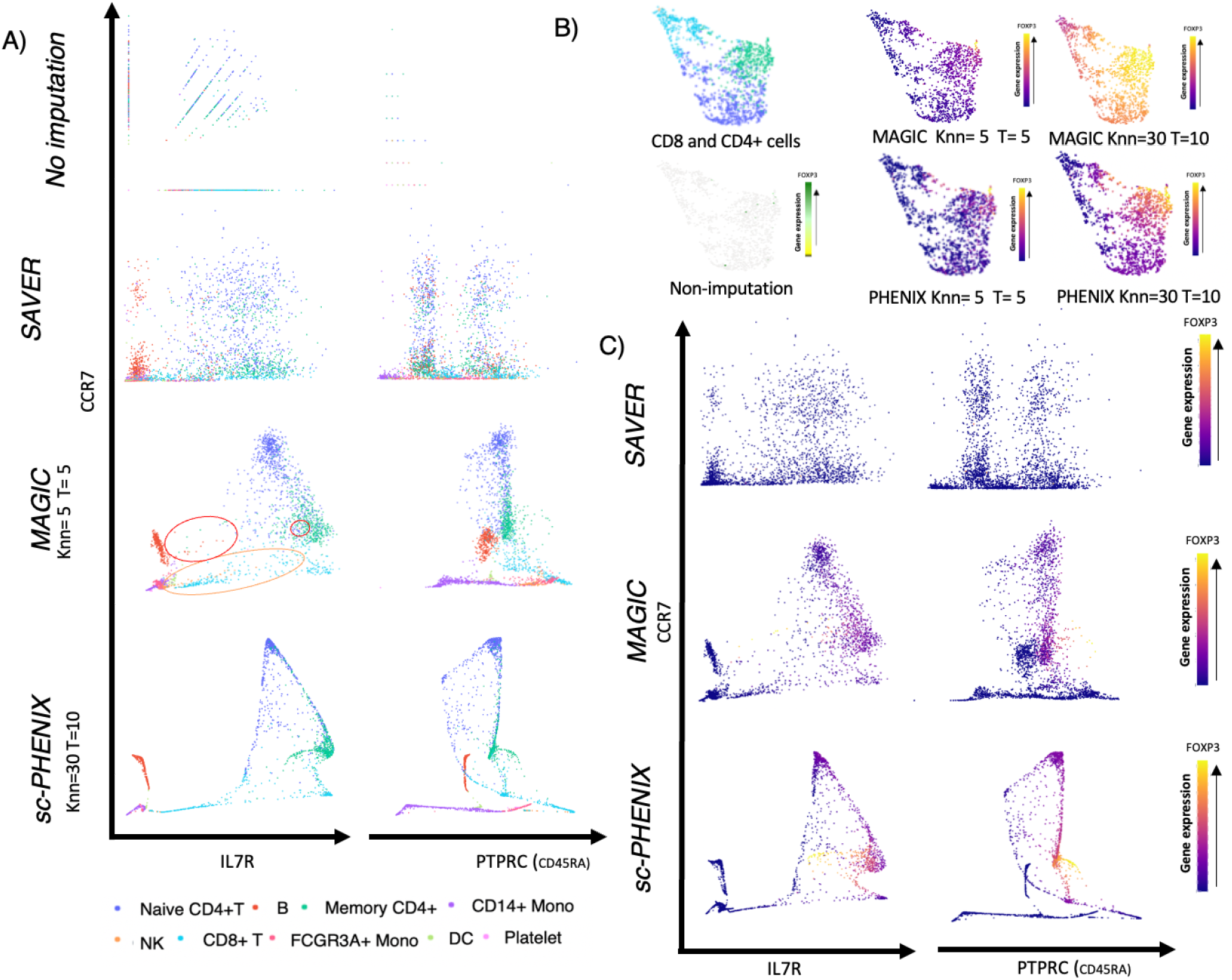
Single-cell RNA-seq imputation algorithms apply to the 3k PBMC dataset. A) IL7R-CCR7 interaction of the non-imputed 3k PBMC dataset and the imputed by MAGIC, SAVER, and sc-PHENIX. Distinct PBMC cell phenotypes are shown below. Red ovals indicate B cells overs-mooted with increasing values of CCR7 in B cells. Orange ovals indicate NK cells over-smoothed with increasing values of CCR7. B) Sections of the UMAP plot of the 3k PBMC dataset showing the CD8+, memory CD4+ and näive CD4+ T cells to show the imputation values of FOXP3 by MAGIC and sc-PHENIX imputation (imputation methods using diffusion maps). The CD8+, memory CD4+ and näive CD4+ T cells clusters (above left). Non-imputed data values of FOXP3 on the UMAP projection (below left). FOXP3 recovered values by MAGIC shown on the UMAP projection, using different values of diffusion parameters of knn and t (above middle and right). FOXP3 recovered values by sc-PHENIX shown on the UMAP projection, using different values of diffusion parameters of knn and t (below middle and right). C) IL7R-CCR7 and PTPRC-CCR7 interactions of the recovered 3k PBMC dataset by MAGIC, SAVER and sc-PHENIX. Recovered values of FOXP3 are shown on the IL7R-CCR7 and PTPRC-CCR7 recovered interactions.

We evaluated the effect of over-smoothing on two gene-gene interactions recovered by sc-PHENIX, MAGIC, and SAVER. Firts, we used the IL7R-CCR7 interaction. The IL7R-CCR7 interaction is related to the transition dynamics of a nävie to a memory CD4+ T-cell state. This transition involves a **CCR7** downregulation [25]. Also, **IL7R** is expressed in both nävie CD4+ and memory CD4+ T cell types but stimulates the proliferation of mature CD4+ T cells [25].

The second gene-gene interaction is CCR7-PTPRC. The expression of CD45RA (also known as **PTPRC**) is generally associated with näive T cells. However, a subset of CD45RA^+^CCR7^-^effector memory T cells re-expresses CD45RA in cytometry experiments[26]. It is interesting to evaluate the recovered expression of the CCR7-ILR7 and CCR7-PTPRC interactions by the scRNA-seq imputation methods. For the non-imputed data (Fig. 4A), it is infrequent to find strong relationships when the data present high rates of dropouts, especially a relationship that explains the transition dynamics of cell states (5). Therefore in the non-imputed data, the IL7R-CCR7 or PTPRC-IL7R interactions are only two examples of a gene-gene interaction misrepresentation.However, MAGIC, sc-PHENIX, and SAVER recover IL7R-CCR7 and CCR7-PTPRC interaction, recreating the biaxial plots typically in flow cytometry. The SAVER (model based imputation method) seems to recover some relationship of the gene-gene interactions but it has no well-defined cluster structure (Fig 4A). Also, there is no present continuum structure showing the transition of naive memory cells. SAVER fails to obtain the gene dynamics of the maturation of the proliferation of mature CD4+ or the CD45RA^+^CCR7^-^effector memory T cells that re-expresses CD45RA^+^.

MAGIC and sc-PHENIX share the same diffusion process (smoothing based methods). Therefore, *knn* and *t* parameters are used in the same way in both methods, the main difference is that MAGIC uses PCA space to measure cell distances and in this section sc-PHENIX uses PCA-UMAP space instead. Moreover, as already mentioned, imputation carries over-smoothing in data. We evaluated the effect of over-smoothing, using distinct increasing values of *knn* and *t* for the IL7R-CCR7 interaction for both diffusion approaches (Fig 4A and B and Section 3). MAGIC recovers IL7R and CCR7 values in other cell phenotypes different from nävie CD4+ and memory CD4+ cell types. Thus, MAGIC over smoothed data (Fig 4A, Fig 5 and, Section 3).

**Fig 5.**
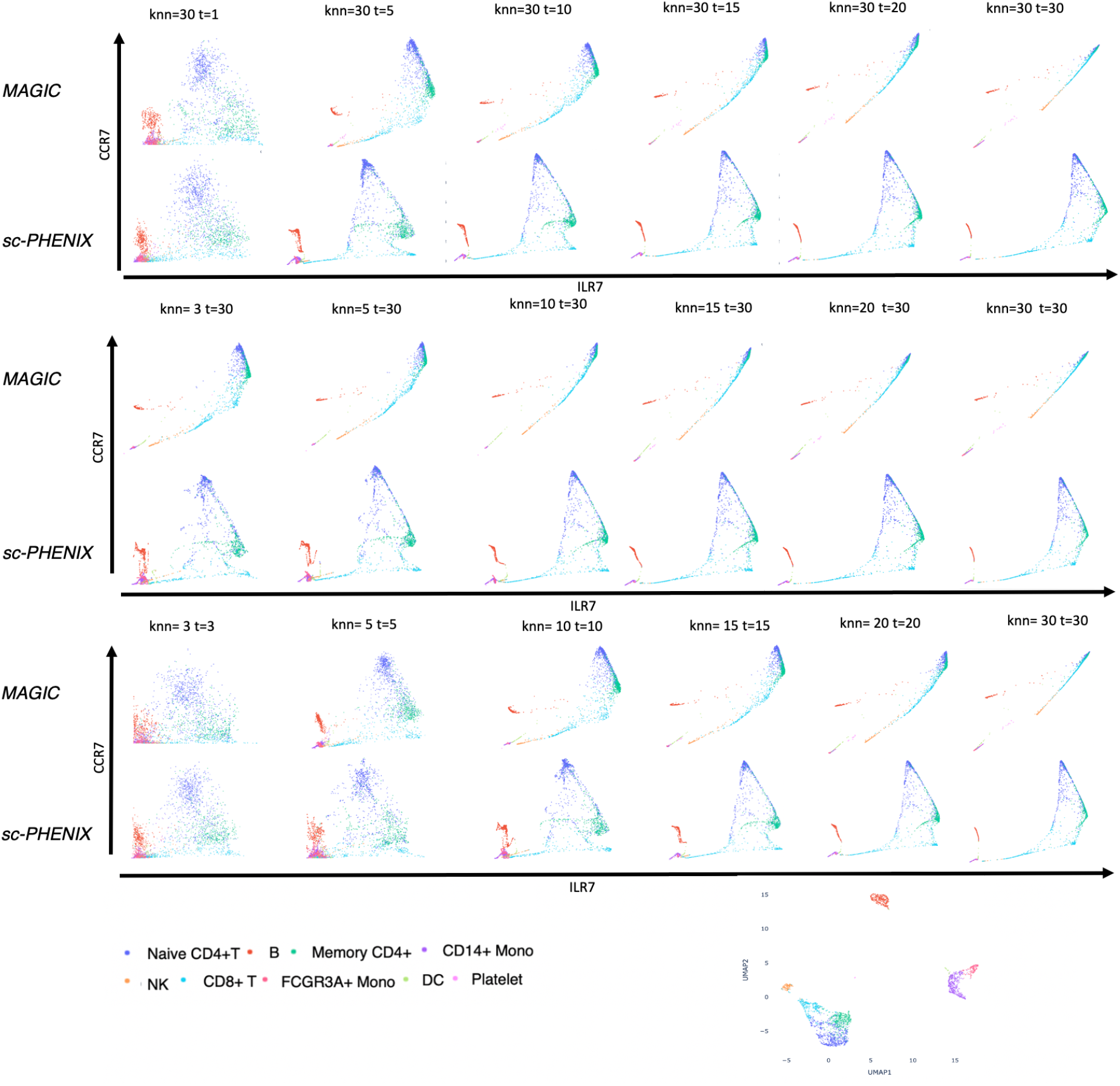
Effect on the gene-gene interactions using different values of the parameter *knn* and *t* used on MAGIC and sc-PHENIX kernels on their imputations. Each dot represents a cell plotted based on their imputed gene expression of IL7R and CCR7 by MAGIC and sc-PHENIX, As *knn* and *t* increases the MAGIC’s recovered IL7R-CCR7 interaction, it starts to recover IL7R and CCR7 values to unrelated cell phenotypes. Also the continuum structure is lost by distorting the dynamic transition of naive to memory CD4+T cells. With sc-PHENIX, the IL7R-CCR7 shape is robust along all combinations of diffusion parameters and continuum structure is captured well by matening the dynamic transition of naive to memory CD4+T cells. In below: UMAP plot of PBMC cell phenotypes.

Additionally, in Section 3 we evaluated ILR7-CCR7 interaction recovered by the MAGIC and sc-PHENIX with UMAP and PCA-UMAP space through increasing principal component dimensions combined with low and high values of diffusion parameters (*knn* and *t*). The sc-PHENIX (PCA-UMAP space) imputation maintains a IL7R-CCR7 interaction through different increasing values of diffusion parameters; over-smoothing has not a higher effect compared to MAGIC (Section 3, Fig 3A and B, Fig 4).

However, MAGIC imputation using low values of diffusion parameters (*knn* and *t*), a more “*local imputation*”, recovers to some extent the transition dynamic of the nävie to a memory CD4+ cell state. But we can observe an over-smoothing of IL7R (in high levels) across some NK cells (Fig 4B, knn=5 and t=5 with MAGIC), NK generally do not express IL7R [27]. With MAGIC, increasing values of diffusion parameters (*knn* and *t*), the dynamic transition is lost by over-smoothing (Fig 4B and S3). As a consequence, the continuum structure is also lost.

In both gene-gene interactions recovered by MAGIC, the local and continuum structure is not well captured as in sc-PHENIX using PCA-UMAP space as initialization. To avoid over-smoothing with MAGIC is to generally pick a *knn* such that it is the smallest value that still results in a connected graph[5]. Also mentions that it is robust to the number of PCA dimensions (10 to 100 PC’s). However, we observed in Section 3 that MAGIC is not robust to the number of PCA dimensions nor to local imputation (low diffusion parameters knn and t). Because, the effect of over-smoothing grows as PCA dimensionality and diffusion parameters values increase. Generally, we observe that MAGIC severely distorts data by over-smoothing.

In Section 3 additionally to evaluate over-something with additional genes, we projected the gene expression of two genes (CD8A and FOXP3) on the IL7R-CCR7 interaction. With MAGIC the CD8A (a marker for CD8+ T cell), CD8A expression is over-smoothed in distinct cell phenotypes that are not CD8+ T cells. Also, with MAGIC set to a local imputation (*knn* =5 and *t=5*) this subpopulation appears but as mentioned before, the data is over-smoothed in a local imputation (Fig 4C and S3). Therefore, even though MAGIC is set to a local imputation does not stop over-smooth the data.

With MAGIC we observe (Fig 4B and C and Section 3) the presence of the subpopulation of CD45RA^+^CCR7^-^effector memory T cells that re-expresses CD45RA^+^. Also this population expresses more FOXP3. FOXP3 is a member of the forkhead transcription factor family. Unlike other members, it is mainly expressed in a subset of CD4+ T-cells that play a suppressive role in the immune system [26]. However, in fig 4C or S3, it is easier to detect this subpopulation sc-PHENIX (PCA-UMAP space) rather than MAGIC or sc-PHENIX (only UMAP space). With all combinations of diffusion parameters sc-PHENIX, using only the UMAP space over-smooths the data (seccion S3), this demonstrates the necessity of PCA as initialization for diffusion on UMAP space. Only UMAP does not capture well-high dimensionality of the data. Many studies have shown that PCA helps UMAP to obtain a more detailed local and global structure [14,15].

In general, MAGIC needs to be used with low *knn* and *t* values, and less principal component dimensions for a less over-smoothed imputation. Contrastingly, sc-PHENIX is more robust to not induce over-smoothing by increasing PCA dimensions, *knn* and *t*. Thus, sc-PHENIX using PCA-UMAP space captures more of the variability and portrays a more accurate graph, without distorting the recovered expression driven by a diffusion on PCA space.

As already mentioned in the previous section the MAGIC (diffusion on PCA space) will over-smoothing scRNA-seq data because of the artifact that renders a connection of densities from distinct cell phenotypes. This makes distinct cell phenotypes similar and imputation recovers similar expressions among these distinct cell phenotypes. We only show in this section a small number of cell phenotypes. However, with MAGIC the over-smoothing will increase as more distinct cell phenotypes are in an experiment due to a more chances of connections among distinct cell phenotypes that are near to each other in the high dimensional PCA space. However, sc-PHENIX mitigates this artifact effect.

### Evaluation of Adult Mouse Visual Cortex Cell Dataset

We use the adult mouse visual cortex cell dataset to evaluate the over-smoothing of sc-PHENIX and MAGIC (both diffusion-based methods) in a more diverse cell phenotype and in a different biological system. This scRNA-seq dataset consists of different sub-cell phenotypes(more than previous 3k PBMC dataset). The main cell phenotypes are GABAergic, glutamatergic and non-neuronal cell phenotypes [24]. In this dataset is already established the DEG (Differential Expressed Genes) for specific phenotypes [24] The idea in this evaluation is that a good imputation will not over-smooth these markers to distinct cell phenotypes. Therefore, we evaluate over-smoothing of specific cell phenotype gene markers using MAGIC and sc-PHENIX. For example, Flt1 marker corresponds to endothelial and smooth muscle cells, *Vip* and *Chat* markers for Vip cells, *Sst* marker for Sst cells and *Serpinb11* marker for Gluta_L6b.

In fig 6, we observe visually the recovered marker expression by MAGIC and sc-PHENIX on UMAP plots. Thus, MAGIC and sc-PHENIX have different outcomes regarding over-smoothing along distinct cell phenotypes using different increasing combinations of PCA dimensions, *knn* and *t*. In fig 6, it is clear the effect of over-smoothing of MAGIC through increasing value parameters and PCA dimensionality.We observe in Fig 6A that increasing *knn, t* and *PCA* dimensions, with MAGIC increases over-smoothing more than sc-PHENIX by not maintaining the expression of any gene marker in their respective cell phenotype cluster (visualized in the UMAP projection).

**Fig 6.**
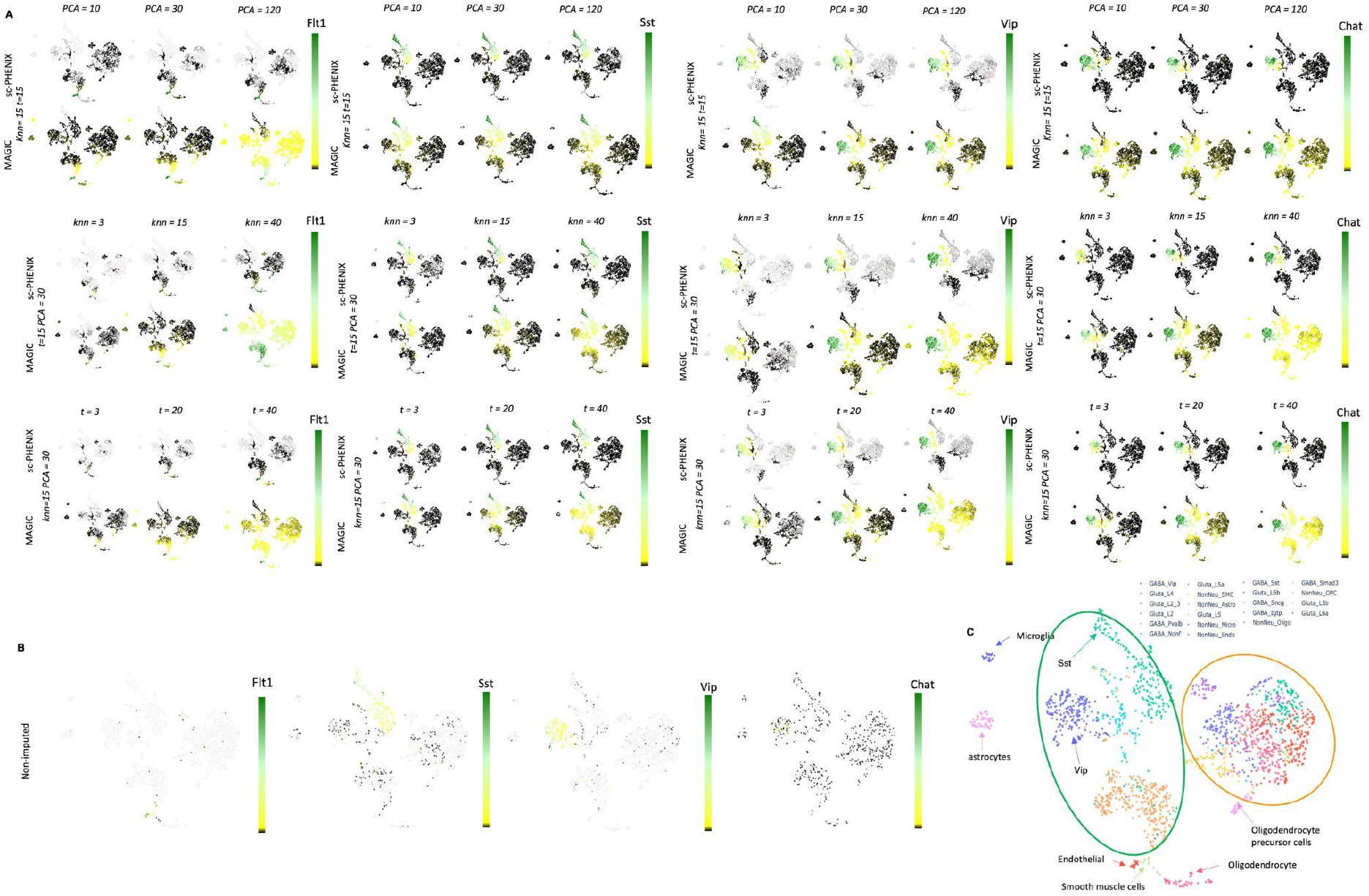
Imputation of the adult mouse visual cortex using MAGIC and sc-PHENIX. A) Recovered Flt, Sst, Vip and CHAT expression are visualized on UMAP projection of the adult mouse visual cortex cells dataset. B) The non-imputed expression values of Flt, Sst, Vip and Chat are visualized on UMAP projection of the adult mouse visual cortex cells dataset. C) The 2D UMAP projection of the adult mouse visual cortex cells dataset, three main clusters are GABAergic (green circle), glutamatergic (orange circle) and non-neuronal cell types. Different parameters were used to see the effect of over-smoothing for MAGIC and sc-PHENIX methods.

In addition, we quantified the imputation performance of both methods to maintain the differential expression of specific gene markers in their respective cell phenotypes (Fig 7). This is a classification problem where there is or is not differential expression of an imputed gene marker on a cell-phenotype and at the same time using more combinations of diffusion parameters (*knn* and *t*) and PCA dimensions. Thus, we can evaluate the precision, recall and f1-score metrics. We used the Flt1 marker for NonNeu_Endo cells and NonNeu_SMC cells; Sst marker for GABA_Sst cells; Chat marker for GABA_Vip cells; Serpin11 for Gluta_L6B cells type. We observed in more combinations of increasing parameters and PCA dimensions that the performance of imputation de decreases more with MAGIC than sc-PHENIX. Thus, indicating that sc-PHENIX in general is more reliable to avoid over-smoothing and deals with the course of dimensionality better than MAGIC(this reflected in the imputation performance and previous sections).

**Fig 7.**
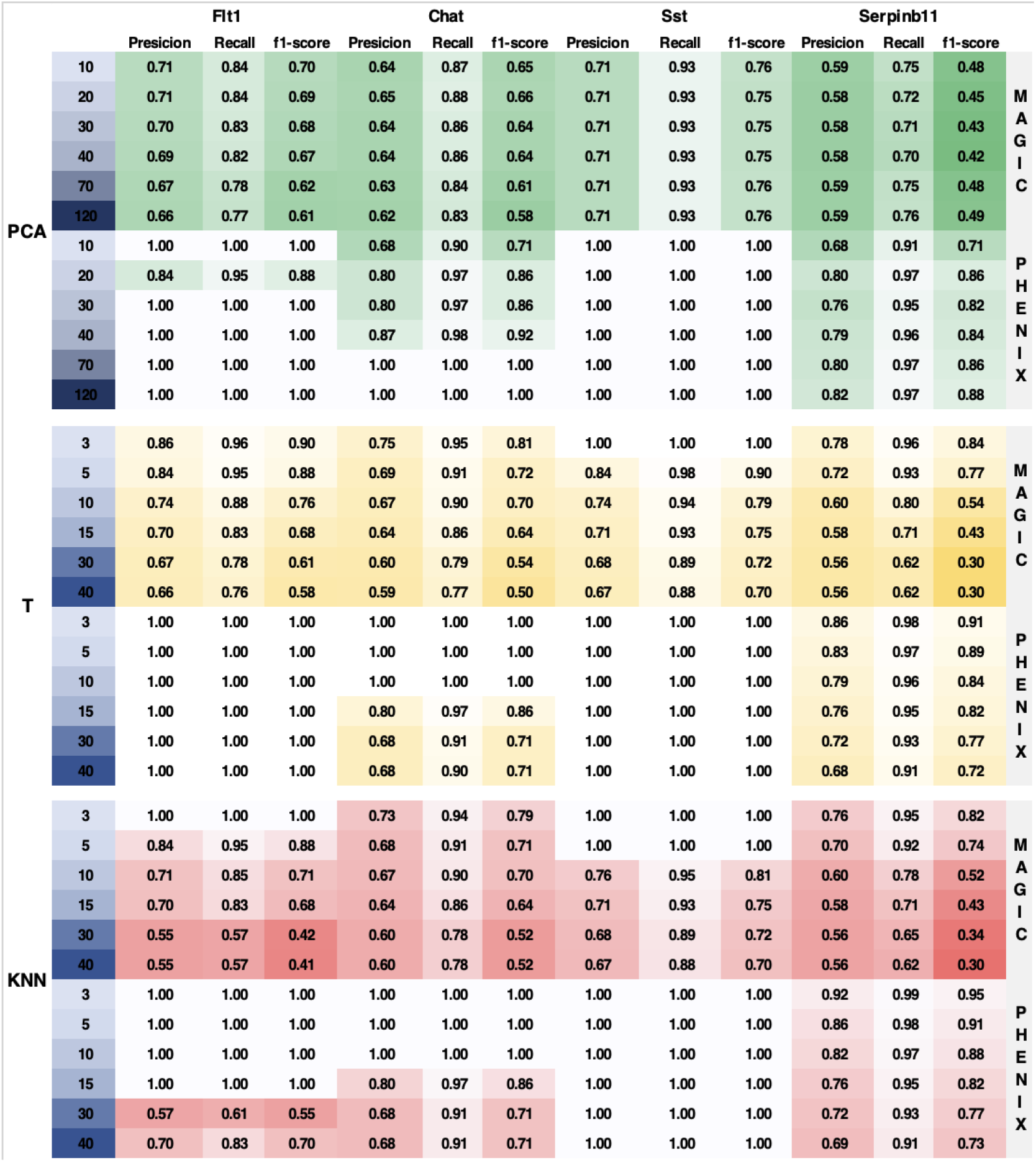
Smoothing performance of MAGIC and sc-PHENIX imputation through different increasing combinations of parameters. Here we show the Precision, Recall and f1-score performance metrics for imputation using different combinations parameters of *knn*, t and PCA dimensions. We used Flt1(NonNeu_Endo and NonNeu_SMC cell types), Chat(GABA_Vip cell type), Sst(GABA_Sst cell type) and Serpin11(Gluta_L6B cell type) Gene markers. The differential expression for the imputed gene markers was set to Fold Change = 2.0 and FWRD = 0.05 using Tukey’s HSD (honestly significant difference). Note: For increasing values for PCA we set it to *knn*=15 and t =15. For increasing values for t we set it to *knn*=15 and PCA= 30. For increasing values of *knn* we set t=15 and PCA= 30 same as Fig 5.

The method sc-PHENIX avoids over-smoothing because the MDS plots of the *M^t^* in previous sections for this scRNA-seq dataset, we observed that sc-PHENIX correctly separates densities of distinct cell phenotypes (GABAenergic, glutamatergic and non neuronal cells) by PCA-UMAP initialization (Fig 2K). In contrast, MAGIC using only PCA initialization does not achieve a well-performance with any combination of parameters; there is over-smoothing of all markers, even with local imputations (low values of *knn* and *t*), see Fig 6 and Fig7. MAGIC over-smooths data because it does not correctly separets densities of distinct cell phenotypes (GABAenergic, glutamatergic and non neuronal cells) in Fig 2I.

Imputation via diffusion on PCA (MAGIC) there is always some inherent danger in over-smoothing the data that has complex cell phenotypes such as Fig 6C. PCA space does not separate well densities from distinct clusters enough to avoid spurious connections by diffusion (already established in a previou section for this dataset, see Fig 2I). Thus, more distinct cell phenotypes increase the chances of over-smoothing with MAGIC.

## Discussion

We present the imputation algorithm sc-PHENIX that captures with a well-balance the local, global and continuum structure of the data avoiding the undesirable effect of over-smoothing. Imputation allows to compensate for the undesirable effects of dropout events that generate distortion on gene distributions. However, the majority of the methods do not improve the performance in downstream analyses compared to no imputation[6]. Also, it is already known that diffusion-based imputation methods retrieve many differentially expressed genes [14], but with our approach, these differential expressed genes are not affected by over-smoothing avoiding spurious gene interactions. Additionally, by sc-PHENIX the gene expression dynamics can be evaluated where model-based methods like SAVER fail (Fig 4A, 4C).

We also provide MDS plots to visualize the exponentialized Markov matrix as a diagnostic plot, visualizing the cell-neighborhood. The MDS plot shows which regions can be at risk of over-smoothing due to spurious neighbors (closeness of distinct phenotypes) as an incorrect association artifact due to a bad choice of parameters. So that the user carefully adjusts the diffusion parameters and the number of PCA dimensions involved in the imputation. Moreover, this can be done only if the user has label information on the cellular phenotypes. Following the sc-PHENIX advantages in the MDS of the *M^t^* plots generated by the sc-PHENIX, we observed better results than PHATE, an algorithm supposedly designed to visualize high-dimensional data; our approach captures the local, global and continuum structure. Therefore, sc-PHENIX computes a more accurate approximation of the real underlying manifold of high-dimensional data (in its *M^t^*). That is better than specialized methods such PHATE or UMAP. In [10] mentions that UMAP does not properly capture the continuum structure of data. However, our MDS of the *M^t^* visualization method transforms the UMAP space(10~100 UMAP dimensions) into a low dimensional manifold that captures the local, global but most importantly the continuum structure. This for imputation or visualization porpurses with sc-PHENIX. The explanation that sc-PHENIX captures well the balance of these data structures is because the markov matrix contamples more than two or three UMAP dimensions. Therefore, the *M^t^* into sc-PHENIX that uses the initialization of PCA-UMAP space, approximates more to the real and underlying manifold of the scRNA-seq data compared to UMAP, PHATE and MAGIC. Visualization is still an area that we will develop in the future.

Here we show that diffusion on PCA space (MAGIC approach) generates an artifact in which densities of different clusters come together becoming spurious near neighbors. Therefore, this generates distortions in the recovered gene expression. Thus, diffusion on PCA space over-smooths data by increasing diffusion process parameters *knn* and *t*. That is not surprising at all if the computed underlying manifold does not approximate the distances from high dimensionality, diffusion on PCA fails in this task. Thus, indicating the limitations of diffusion on PCA space against the effect of the curse of dimensionality, this is a ubiquitous problem in machine learning. The sc-PHENIX (diffusion on PCA-UMAP space) is more robust against over-smoothing caused by increasing PCA dimensions. Therefore, our approach contemplets more variability to capture the underlying manifold where scRNA-seq lies.

Now, sc-PHENIX use UMAP, therefore can use its different variants such as: supervised UMAP (embeddings for highly heterogeneous data), non-parametric UMAP (embedding the connectivity matrix with neural networks), combination of UMAP models (embedding integrating different datasets), Mutual k-NN Graph (Improving the Separation Between Similar Classes) and DensMAP (Better Preserving Local Density). For example, UMAP can combine two topologies from two omics technologies dataset such as CITE-seq (single-cell transcriptomics and proteomics profiles in one experiment). Then having combined these two topologies in an embedding, sc-PHENIX can use it to impute the single-cell-transcriptomic matrix or the proteomic. Another possible implementation is the high-heterogeneous microbiota data from different technologies (16S rRNA and shotgun Whole Metagenome Sequence), the microbiota matrix contains many zeros and it drives distortion of the microbe abundance distributions and few differential microbes among studie cases. It is impossible to obtain a well-cluster structure among samples (sparsity aggravates this), especially in clinical data among patients. The manifold of the supervised UMAP embedding will lead sc-PHENIX imputation to obtain more differential microbes among studie cases(work in progress). Diffusion via PCA space(MAGIC) can render a global picture of the recovered imputed data better than other current imputation methods. However, our novel method renders finer local, global and continuum structure considerations for imputation. Furthermore, gene set enrichment analysis(GSEA) with the imputed data with sc-PHENIX shows better results in a biological context compared to other imputation methods (data not shown, work in progress).

## Methods

### Overview of the sc-PHENIX

sc-PHENIX, is a computational method for scRNA-seq data imputation, uses an input of *n*-by-*m* count matrix *D* and returns an imputed count matrix D_imputed_. The D_imputed_ matrix represents the likely expression for each single cell, based on data diffusion between similar cells (Fig 1).

The key of a correct imputation using sc-PHENIX is due to a well-constructed affinity matrix *M*. So the construction of *M* is mentioned below:

1. The count matrix *D* is transformed using PCA to obtain a lower-dimensional representation *D_PCA_*. 2) Then, the low-dimensional representation *D_PCA_* is transformed using UMAP *B_PCA–UMAP_*. 3) Computation of a cell-cell distance matrix *Distance_matrix*. 4) Computation of the affinity matrix *A* based on *Distance_matrix*, via an adaptive Gaussian kernel function. 5) Symmetrization of *A* using an additive approach. 6) Row-stochastic Markov-normalization of *A* into Markov matrix *M*.

The Markov affinity matrix *M* represents the probability distribution of transitioning from one cell to another (the probability of cell “*i*” and cell “*j*” are similar).

After the construction of the Markov affinity matrix *M*, data diffusion is through exponentiation *t* of *M*. The effect of *t* exponentiation filters out noise and increases the similarity based on strong trends in the data. Thus, increasing the weight (probability) among phenotypically similar cells. In contrast, phenotypically distinct cells are down-weighted. This noise is presented by technical errors such as dropouts, artifacts, and a small and randomized ratio of the capture of transcripts. Noise does not have a strong trend in the manifold, but similar cells do have.

So this filtration step is crucial for the elimination of noise. Once the Affinity matrix M is exponentiated to recover the lost values in the scRNA-seq data via Matrix multiplication.

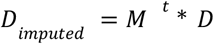

There is the option to rescale the values so that the max value for each gene equals the 99^th^ percentile of the original data (same as older versions of MAGIC).

Also, there is the option of using other manifold representations, not just PCA-UMAP space, any other machine learning that generates a lower-dimensional and denoised representation of which the data could lie can be used as an input for sc-PHENIX to create the *Distance _matrix*. If PCA space is used sc-PHENIX will render the same results as MAGIC.

#### Preprocessing of the row count single-cell matrix

Before applying sc-PHENIX is recommended to do quality control [5]. Thus, filtering by removing low-quality cells or removing empty droplets (for droplet-based technologies), removing empty genes, etc. Then, apply a library size normalization to finally obtain the scRNA-seq input data for sc-PHENIX imputation.

Doing a library size normalization (in a technology-dependent manner), effectively eliminates cell size as a signal in the measurement for the purposes of constructing the affinity matrix and thus the resulting weighted neighborhood is not biased by cell size.

#### PCA-UMAP manifold and creation of the Distance cells Matrix

There are two start points to create the the distance cell matrix (Fig 1) that would have significant differences on the imputation results:

1. The user has the option to bring their denoise representation of the scRNA-seq original data in a lower dimension to be imputed that can be interpreted as Euclidean or any other simple metric distance (cosine, Manhattan, etc.); using the most variable genes or any other dimensionality reduction machine learning technique, etc. Here the user can include PCA (just like MAGIC) or UMAP.
2. The preprocessed single-cell count matrix is reduced dimensionality by using PCA as an informative initialization step for UMAP to obtain the PCA-UMAP space. UMAP embeddings are projections that can be interpreted as Euclidean distance.

The main difference between sc-PHENIX and MAGIC lies in the use of de PCA-UMAP space to calculate the distance matrix of cells rather than using just PCA space like MAGIC.

It is already known that a PCA-UMAP manifold is necessary to reveal fine-scale (local) structure that only PCA or UMAP can not achieve, especially for noisy and high-dimensional data [12]. Principal Components Analysis (PCA) is a critical initialization step for UMAP for helping to preserve global [11] and local data structure [12]. Thus, creating a PCA-UMAP space of which lies the scRNA-seq manifold more accurately. This lower dimension representation of the high-dimensional scRNA-seq data is the key to better imputation performance when sc-PHENIX is used. However the UMAP method using a very low dimensional representation (2D or 3D) of UMAP is not recommended, as high-dimensional distances cannot be fully preserved in lower dimensions especially for scRNA-seq data [28].

The advantage of UMAP is that there is really no requirement to stop at n_components=3 (UMAP dimension). If you are interested in (density based) clustering or other machine learning techniques (such as sc-PHENIX), it can be beneficial to pick a larger embedding dimension (say 10, or 50) closer to the dimension of the underlying manifold on which sc-RNA data lies.

#### Creation of the Affinity Matrix

In order to create the Affinity Matrix A, sc-PHENIX uses an adaptive gaussian kernel. The advantages of using an adaptive kernel are already known for stability of the imputation [5]. We use the same adaptive kernel used in MAGIC, defined as:

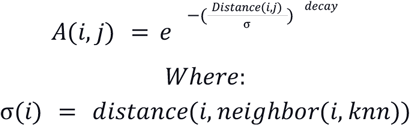

And the reason of the success of the adaptive kernel is due to the adaptability of the bandwidth σ in context of his neighbors, the value σ(*i*) for a cell *i* is the distance to its kath-nearest neighbors (*knn*).

After applying the adaptive kernel to the distance matrix, a non-symmetric affinity matrix A is generated. The adaptive kernel results in an asymmetric affinity matrix where *A*(*i, j*) ≠ *A*(*j, i*), and which it is needed to symmetrize to achieve these desired properties for A. Symmetrization of the affinity matrix is as follows:

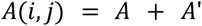

#### Row-stochastic Markov-normalization of *A* into Markov matrix *M*

A row normalization of A is need it to create the Markov transition matrix *M*. Each row of this Markov transition matrix must sum 1 showing the probability distribution of the transition from one cell to every other cell in the scRNA-seq data. Row normalization is calculated by dividing each entry in A by the sum of row affinities:

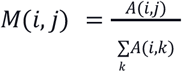

#### Data diffusion and imputation

Once *M* is calculated, the diffusion process starts with the exponentiation of *M*, this means that phenotypically similar cells should have strongly weighted affinities and noise/spurious neighbors are down-weighted in this diffusion process. Raising the *M* to the power *t* generates this *M^t^* (*i, j*) of which represents the probability that a random walk of length t starting at cell i will reach cell j, same as MAGIC we call *t* the ‘*diffusion time*’ In the first few steps of *diffusion time*, sc PHENIX learns the manifold structure and removes the noise dimensions.

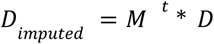

There is the option of rescaling the data, the max value for each gene equals the 99th percentile of the original data, same as older versions in MAGIC. This is not recommended to do it, if the single cell matrix has negative values[5].

### Multidimensional scaling of exponiated transition Markov matrix

The PCA space and the exponentiated transition Markov matrix(from PCA and PCA-UMAP initialization) is transformed into a dissimilar matrix (Distance matrix) to apply a Multidimensional scaling. For a faster optimization and a more accurate preservation of the high dimensionality distances in the low dimensional manifold we used the algorithm in this multidimensional scaling approach[29] to minimizes its energy function, known as stress, by using stochastic gradient descent to move a single pair of vertices at a time. Their results show that stochastic gradient descent can reach lower stress levels faster and more consistently than majorization, without needing help from a good initialization. The approach in [29] can project data from a distance matrix into 2D and 3D dimensions(MDS dimensions).

### scRNA-seq datasets

3k PBMCs(Peripheral Blood Mononuclear Cells) from a Healthy Donor, PBMCs are primary cells with relatively small amounts of RNA (~1pg RNA/cell). 2,700 cells detected: https://support.10xgenomics.com/single-cell-gene-expression/datasets/1.1.0/pbmc3k

Quality Control, normalize (libsize) and transform (Log +1) same as Seurat method in R: https://satijalab.org/seurat/articles/pbmc3k_tutorial.html

Adult mouse visual cortex (RPKM values for 24,057 genes and 1,679 cells) https://singlecell.broadinstitute.org/single_cell/study/SCP6/a-transcriptomic-taxonomy-of-adult-mouse-visual-cortex-visp#study-download

MAGIC EMT (7523 cells and 28910 genes,: https://github.com/KrishnaswamyLab/MAGIC/blob/master/data/HMLE_TGFb_day_8_10.csv.gz, GEO Accession: GSE114397.

## Acknowledgements

OR thanks the financial support from CONACYT (Grant Ciencia de Frontera 2019, FORDECYT-PRONACES/425859/2020), PAPIIT-UNAM (IA202720), and an internal grant from the National Institute of Genomic Medicine (INMEGEN, México). PM-C is a doctoral student from Programa de Doctorado en Ciencias Biomédicas, Universidad Nacional Autónoma de México (UNAM) and received fellowship to CVU 855825 from CONACYT, México. This paper is part of the doctoral thesis and the requirements to obtain the degree of Doctor in Science to PM-C. YEM-L is a doctoral student from Programa de Doctorado en Ciencias Médicas, Odontológicas y de la Salud by Universidad Nacional Autónoma de México (UNAM) and received fellowship to CVU 629384 from CONACYT. DAE-H is a postdoctoral research associate at INMEGEN and received CONACyT fellowship to CVU 420693. DN-R is a student from the Master in Sciences program in Ciencias Bioquímicas, UNAM and received CONACyT fellowship to CVU 1083211. JPS-C is a student from the Master in Sciences program in Ciencias Bioquímicas, UNAM and received CONACyT fellowship to CVU 1005702. DG-V is a student from the Master in Sciences program in Ciencias Bioquímicas, UNAM and received CONACyT fellowship to CVU 1083058.

## Supporting information

S1 Section. sc-PHENIX imputation using PCA initialization to reproduce the same results as MAGIC.

(PDF)

https://github.com/resendislab/sc-PHENIX/blob/main/Sections/Section%20S1.pdf

S2 Section. Corrupted image digits data visualized using PCA, UMAP and PHATE through different rates of induced dropout-events. The UMAP and PHATE had an initialization of 50 PCs.

(PDF)

https://github.com/resendislab/sc-PHENIX/blob/main/Sections/Section%20%20S2.pdf

S3 Section. Recovered CCR7-IL7R interaction by sc PHENIX and MAGIC.

We used different combinations of diffusion parameters (knn, t and PCA dimensions) to observe the over-smoothing of both methods.

https://github.com/resendislab/sc-PHENIX/blob/main/Sections/Section%20S3.pdf

S1 Fig. Interactive 3D MDS plot of the PCA space. (500 PC,500 PC’s, MNIST).

(HTML)

https://github.com/resendislab/PHENIX/blob/main/Supplementary%20figures/FIG%20S1%203D%20MSD%20plot%20500%20PC.html

S2 Fig. Interactive 3D MDS plot of the exponentiated Markov matrix of diffusion on PCA space(500 PC’s input, MNIST).

(HTML)

https://github.com/resendislab/PHENIX/blob/main/Supplementary%20figures/FIG%20S2%203D%20PCA_DIFF_MSD%20plot%20500%20PC.html

S3 Fig. Interactive 3D MDS plot of the exponentiated Markov matrix of diffusion on PCA-UMAP space(500 PC’s transformed into 60 UMAP components as input, MNIST).

(HTML)

https://github.com/resendislab/PHENIX/blob/main/Supplementary%20figures/FIG%20S3%203D%20PCA_UMAP_DIFF_MSD%20plot%20500%20PC.html

S4 Fig. 2D PHATE plot of MNIST dataset(knn=5, n_pca=500 PHATE parameters).

We used a knn= 5 for PHATE plots, getting a higher knn cluster tends to lose local structure.

https://github.com/resendislab/PHENIX/blob/main/Supplementary%20figures/Fig%20S4%20phate.png

(PNG)

S5 Fig. 2D PHATE plot of MNIST dataset merge with MNIST images

We used a knn= 5 for PHATE plots, getting a higher knn cluster tends to lose local structure.

https://github.com/resendislab/PHENIX/blob/main/Supplementary%20figures/Fig%20S5%20phate%20image%20MNIST%20numbers.png

(PNG)

S6 Fig. 2D PHATE plot 500 PC neuronal

We used a knn= 5 for PHATE plots, getting a higher knn cluster tends to lose local structure.

https://github.com/resendislab/PHENIX/blob/main/Supplementary%20figures/Fig%20S7%202D%20phate%20plot%20500%20PC%20neuronal.png

(PNG)

S7 Fig. 2D PCA_DIFF_MSD plot (500 PC) merged with MNIST images

https://github.com/resendislab/PHENIX/blob/main/Supplementary%20figures/Fig%20S8%202D%20PCA_DIFF_MSD%20plot%20500%20PC.png

(PNG)

S8 Fig. 2D PCA_UMAP_DIFF_MSD plot (500 PC) merged with MNIST images

https://github.com/resendislab/PHENIX/blob/main/Supplementary%20figures/Fig%20S9%202D%20PCA_UMAP_DIFF_MSD%20plot%20500%20PC%20.png

(PNG)

## Notes

https://github.com/resendislab/sc-PHENIX

### Competing Interest Statement

The authors have declared no competing interest.

### Summary of Updates

The only change was is the Correspondence author that is Resendis-Antonio Osbaldo. This in the PDF

## References

1. AlJanahi AA, Danielsen M, Dunbar CE. An Introduction to the Analysis of Single-Cell RNA-Sequencing Data. Molecular Therapy - Methods & Clinical Development. 2018 Sep;10:189–96.

2. Kharchenko PV, Silberstein L, Scadden DT. Bayesian approach to single-cell differential expression analysis. Nature Methods. 2014 May 18;11(7):740–2.

3. Stegle O, Teichmann SA, Marioni JC. Computational and analytical challenges in single-cell transcriptomics. Nature Reviews Genetics. 2015 Jan 28;16(3):133–45.

4. Grün D, Kester L, van Oudenaarden A. Validation of noise models for single-cell transcriptomics. Nature Methods. 2014 Apr 20;11(6):637–40.

5. van Dijk D, Sharma R, Nainys J, Yim K, Kathail P, Carr AJ, et al. Recovering Gene Interactions from Single-Cell Data Using Data Diffusion. Cell. 2018 Jul;174(3):716–729.e27.

6. Hou W, Ji Z, Ji H, Hicks SC. A Systematic Evaluation of Single-cell RNA-sequencing Imputation Methods [Internet]. Cold Spring Harbor Laboratory; 2020 Jan [cited 2022 Jan 26]. Available from: http://dx.doi.org/10.1101/2020.01.29.925974

7. Rostom R, Svensson V, Teichmann SA, Kar G. Computational approaches for interpreting scRNA-seq data. FEBS Letters. 2017 Jun 12;591(15):2213–25.

8. Kumari S, Jayaram B. Measuring Concentration of Distances—An Effective and Efficient Empirical Index. IEEE Transactions on Knowledge and Data Engineering. 2017 Feb 1;29(2):373–86.

9. Andrews TS, Hemberg M. Identifying cell populations with scRNASeq. Molecular Aspects of Medicine. 2018 Feb;59:114–22.

10. Peres-Neto PR, Jackson DA, Somers KM. How many principal components? stopping rules for determining the number of non-trivial axes revisited. Computational Statistics & Data Analysis. 2005 Jun;49(4):974–97.

11. Amir ED, Davis KL, Tadmor MD, Simonds EF, Levine JH, Bendall SC, et al. viSNE enables visualization of high dimensional single-cell data and reveals phenotypic heterogeneity of leukemia. Nature Biotechnology. 2013 May 19;31(6):545–52.

12. McInnes L, Healy J, Saul N, Großberger L. UMAP: Uniform Manifold Approximation and Projection. Journal of Open Source Software. 2018 Sep 2;3(29):861.

13. Moon KR, van Dijk D, Wang Z, Gigante S, Burkhardt DB, Chen WS, et al. Visualizing structure and transitions in high-dimensional biological data. Nature Biotechnology. 2019 Dec;37(12):1482–92.

14. Kobak D, Linderman GC. Initialization is critical for preserving global data structure in both t-SNE and UMAP. Nature Biotechnology. 2021 Feb;39(2):156–7.

15. Sakaue S, Hirata J, Kanai M, Suzuki K, Akiyama M, Lai Too C, et al. Dimensionality reduction reveals fine-scale structure in the Japanese population with consequences for polygenic risk prediction. Nature Communications. 2020 Mar 26;11(1).

16. Malzer C, Baum M. A Hybrid Approach To Hierarchical Density-based Cluster Selection. In: 2020 IEEE International Conference on Multisensor Fusion and Integration for Intelligent Systems (MFI) [Internet]. IEEE; 2020 [cited 2022 Jan 26]. Available from: http://dx.doi.org/10.1109/mfi49285.2020.9235263

17. Tjärnberg A, Mahmood O, Jackson CA, Saldi G-A, Cho K, Christiaen LA, et al. Optimal tuning of weighted kNN- and diffusion-based methods for denoising single cell genomics data. PLOS Computational Biology. 2021 Jan 7;17(1):e1008569.

18. Patruno L, Maspero D, Craighero F, Angaroni F, Antoniotti M, Graudenzi A. A review of computational strategies for denoising and imputation of single-cell transcriptomic data. Briefings in Bioinformatics. 2020 Oct 1;

19. Huang M, Wang J, Torre E, Dueck H, Shaffer S, Bonasio R, et al. SAVER: Gene expression recovery for UMI-based single cell RNA sequencing [Internet]. Cold Spring Harbor Laboratory; 2017 May [cited 2022 Jan 27]. Available from: http://dx.doi.org/10.1101/138677

20. Coifman RR, Lafon S, Lee AB, Maggioni M, Nadler B, Warner F, et al. Geometric diffusions as a tool for harmonic analysis and structure definition of data: diffusion maps. Proceedings of the National Academy of Sciences of the United States of America. 2005 May 24;102(21):7426–31.

21. Elyanow R, Dumitrascu B, Engelhardt BE, Raphael BJ. netNMF-sc: Leveraging gene-gene interactions for imputation and dimensionality reduction in single-cell expression analysis [Internet]. Cold Spring Harbor Laboratory; 2019 Feb [cited 2022 Jan 26]. Available from: http://dx.doi.org/10.1101/544346

22. Haghverdi L, Buettner F, Theis FJ. Diffusion maps for high-dimensional single-cell analysis of differentiation data. Bioinformatics. 2015 May 21;31(18):2989–98.

23. Li Deng. The MNIST Database of Handwritten Digit Images for Machine Learning Research [Best of the Web]. IEEE Signal Processing Magazine. 2012 Nov;29(6):141–2.

24. Tasic B, Menon V, Nguyen TN, Kim TK, Jarsky T, Yao Z, et al. Adult mouse cortical cell taxonomy revealed by single cell transcriptomics. Nature neuroscience. 2016 Feb;19(2):335–46.

25. Lee MS, Hanspers K, Barker CS, Korn AP, McCune JM. Gene expression profiles during human CD4+ T cell differentiation. International Immunology. 2004 Jun 21;16(8):1109–24.

26. Tian, Babor, Lane, Schulten, Patil, Seumois, et al. Unique phenotypes and clonal expansions of human CD4 effector memory T cells re-expressing CD45RA. Nature Communications. 2017 Nov 13;8(1):1–13.

27. Sheikh A, Abraham N. Interleukin-7 Receptor Alpha in Innate Lymphoid Cells: More Than a Marker. Frontiers in Immunology. 2019 Jan 1;0.

28. Lange, Bergen, Klein, Setty, Reuter, Bakhti, et al. CellRank for directed single-cell fate mapping. Nature Methods. 2022 Jan 13;19(2):159–70.

29. Zheng JX, Pawar S, Goodman DFM. Graph Drawing by Stochastic Gradient Descent. IEEE Transactions on Visualization and Computer Graphics. 2019 Sep 1;25(9):2738–48.

